# Host autophagy mediates organ wasting and nutrient mobilization for tumor growth

**DOI:** 10.1101/2020.12.04.411561

**Authors:** Rojyar Khezri, Petter Holland, Todd Andrew Schoborg, Ifat Abramovich, Szabolcs Takats, Caroline Dillard, Ashish Jain, Fergal OFarrell, Sebastian Wolfgang Schultz, William M. Hagopian, Eduardo Martin Quintana, Rachel Ng, Nadja Sandra Katheder, Mohammed Mahidur Rahman, Andreas Brech, Heinrich Jasper, Nasser M. Rusan, Anne Hope Jahren, Eyal Gottlieb, Tor Erik Rusten

## Abstract

During tumor growth - when nutrient and anabolic demands are high – autophagy supports tumor metabolism and growth through lysosomal organelle turnover and nutrient recycling^1^. Ras-driven tumors additionally invoke non-autonomous autophagy in the microenvironment to support tumor growth, in part through transfer of amino acids^2–4^. Here we uncover a third critical role of autophagy in mediating systemic organ wasting and nutrient mobilization for tumor growth using a well-characterized malignant tumor model in *Drosophila melanogaster*. Micro-computed X-ray tomography and metabolic profiling reveal that *Ras^V12^; scrib*^-/-^ tumors grow 10-fold in volume, while systemic organ wasting unfolds with progressive muscle atrophy, loss of body mass, −motility, −feeding and eventually death. Tissue wasting is found to be mediated by autophagy and results in host mobilization of amino acids and sugars into circulation. Natural abundance Carbon 13 tracing demonstrates that tumor biomass is increasingly derived from host tissues as a nutrient source as wasting progresses. We conclude that host autophagy mediates organ wasting and nutrient mobilization that is utilized for tumor growth.

## Introduction

### Gradual organ atrophy and weight loss ensues during malignant tumor growth

To assess the dynamics of organ atrophy at the whole-animal level, we adopted computed tomography (CT), the gold standard for evaluating adipose and muscle atrophy in cancer patients. Genetically induced GFP-labeled malignant *Ras^V12^; scrib*^-/-^ eye tumors grow and invade the neighboring Central Nervous System (CNS), extend the larval stage and kill the host by day 10-12^5,6^ We optimized a fixation and staining protocol for high resolution micro-CT (μ-CT) imaging of developmentally staged larvae^7^. This enabled ready identification, segmentation and calculation of tumor and organ volumes (Fig. 1a-c Supplementary Video 1-7)^7^ *Ras^V12^; scrib*^-/-^ tumors grow 10-fold in volume while invading and enveloping the brain from day 6 to 10 (Fig. 1c, Supplementary Video 7, quantified in 2a). Conversely, total larval muscle volume is initially similar to control animals carrying benign *Ras^V12^* tumors, and progressively shrink by approximately 50% from day 6 to day 10 (Fig. 1b, quantified in 2b). The fat body, which perform adipose and liver functions, displayed a striking increase in translucency and lipid droplet size (Fig. 2e, Fig 3c) in *Ras; scrib*^-/-^ tumor-carrying animals from day 8^8^, although fat body volume remained unaltered (Fig. 2d). Organ wasting and tumor growth was accompanied by approximately 35% dry weight loss and a progressive loss of motility and feeding from day 8 (Fig. 2f-h). We established that muscle and fat body changes can be imaged and quantified in intact heat-killed whole larvae using a myosin heavy chain-GFP muscle reporter and backlight microscopy (Fig 3a-c, Expanded View Fig. 1a-c). *Ras^V12^; scrib*^-/-^ tumors inhibit ecdysone synthesis and offset the normal pupation at day 6 due to dilp8 secretion from tumor tissue^9,10^. No muscle or adipose tissue atrophy was observed in day 10 animals where secretion of the molting hormone ecdysone was specifically obliterated in *ecdysoneless* (*ecd^1-/-^*) mutants, or by genetic elimination of the ecdysone-producing prothoracic gland cells (Expanded View Fig. 1d-f). Thus, the onset of muscle and adipose tissue wasting *Ras^V12^, scrib*^-/-^ larvae precede the reduction of feeding and reduced motility, arguing together that the wasting responses are not a simple function of reduced food intake or extended larval stage.

**Figure 1.**
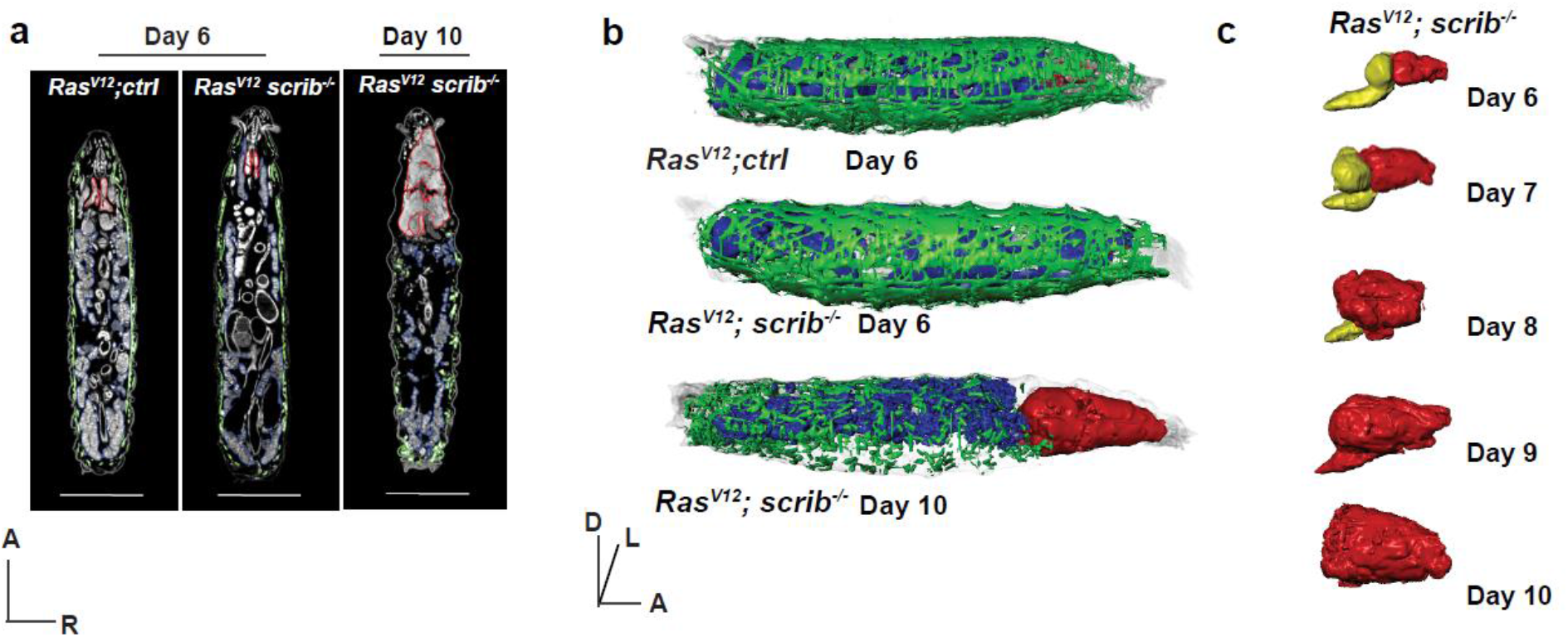
Tumor induced organ wasting. **(a)** Representative 2D μ-CT scans of *Ras^V12^; ctrl* at day 6, *Ras^V12^; scrib*^-/-^ larvae at day 6 and day 10. Muscle (green), fat body (blue) and eye-antennal discs/tumor (red) are outlined. **(b)** Representative 3D rendering of genotypes indicated in (a). Anterior (A), Dorsal (D), Left (L), Right (R). **(c)** Representative 3D rendering of *Ras^V12^; scrib*^-/-^ tumors (red) and central nervous system (yellow), over time.

**Figure 2.**
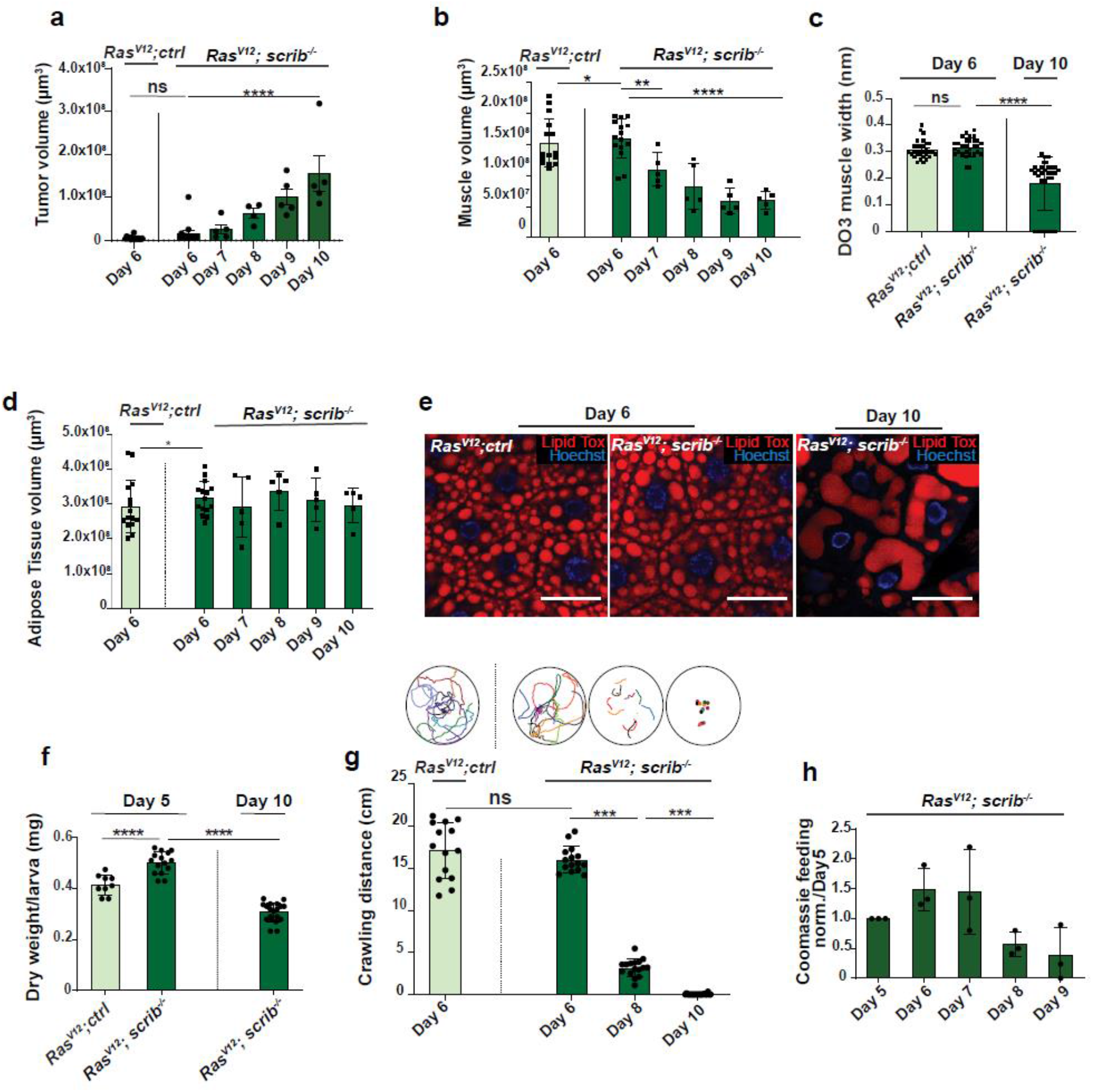
Tumor induced organ wasting. **(a)** Quantification of tumor growth of *Ras^V12^; ctrl* day 6 (n=15) *Ras^V12^; scrib*^-/-^ tumors, day 6 (n=15), day 7 (n=5), day 8 (n=4), day 9 (n=5) and day 10 (n=5). **(b)** Quantifications of muscle volume, of larvae carrying tumors of *Ras^V12^; ctrl* (n=15), and *Ras^V12^; scrib*^-/-^ at day 6(n=15), day 7 (n=5), day 8 (n=5), day 9 (n=5) and dat 10 (n=5). **(c)** Quantification of width of Dorsal Oblique 3 (DO3) muscle in larvae carrying tumors of *Ras^V12^; ctrl* at day 6 (n=29) and *Ras^V12^; scrib*^-/-^at days 6 (n=29) and day 10 (n=31). **(d)** Quantification of adipose tissue volume in larvae carrying tumors of *Ras^V12^; ctrl* at day 6 and *Ras^V12^; scrib*^-/-^at days 6 and day 10. **(e)** Representative confocal images of adipose tissue in *Ras^V12^; ctrl* and *Ras^V12^; scrib*^-/-^ tumors bearing animals at indicated ages. Lipid droplets are highlighted with Lipid Tox staining. **(f)** Quantification of dry weight of *Ras^V12^; ctrl* (n=9), *Ras^V12^; scrib*^-/-^ day 6(n=15) and day 10(n=18) tumor bearing larvae excluding the tumor weight. **(g)** Quantification of larval motility measure by crawling distance and crawling pattern in *Ras^V12^; ctrl* (n=14), *Ras^V12^; scrib*^-/-^ day 6 (n=15), day 8 (n=16) and day 10(n=14), each colored line represents a single larva. **(h)** Coomassie feeding assay to asses food intake of *Ras^V12^; scrib*^-/-^ over time, three repeated measurements of an average food intake of 20 larvae. Values depict mean+s.e.m. of minimum three independent pooled experiments. ns, not significant, \**P* ≤ 0.05, \*\**P* ≤ 0.01, \*\*\**P* ≤ 0.0001 and \*\*\*\**P* < 0.0001, from unpaired, two-tailed test. Scale bar: (1a): 1mm, (1h):50 μm.

**Figure 3.**
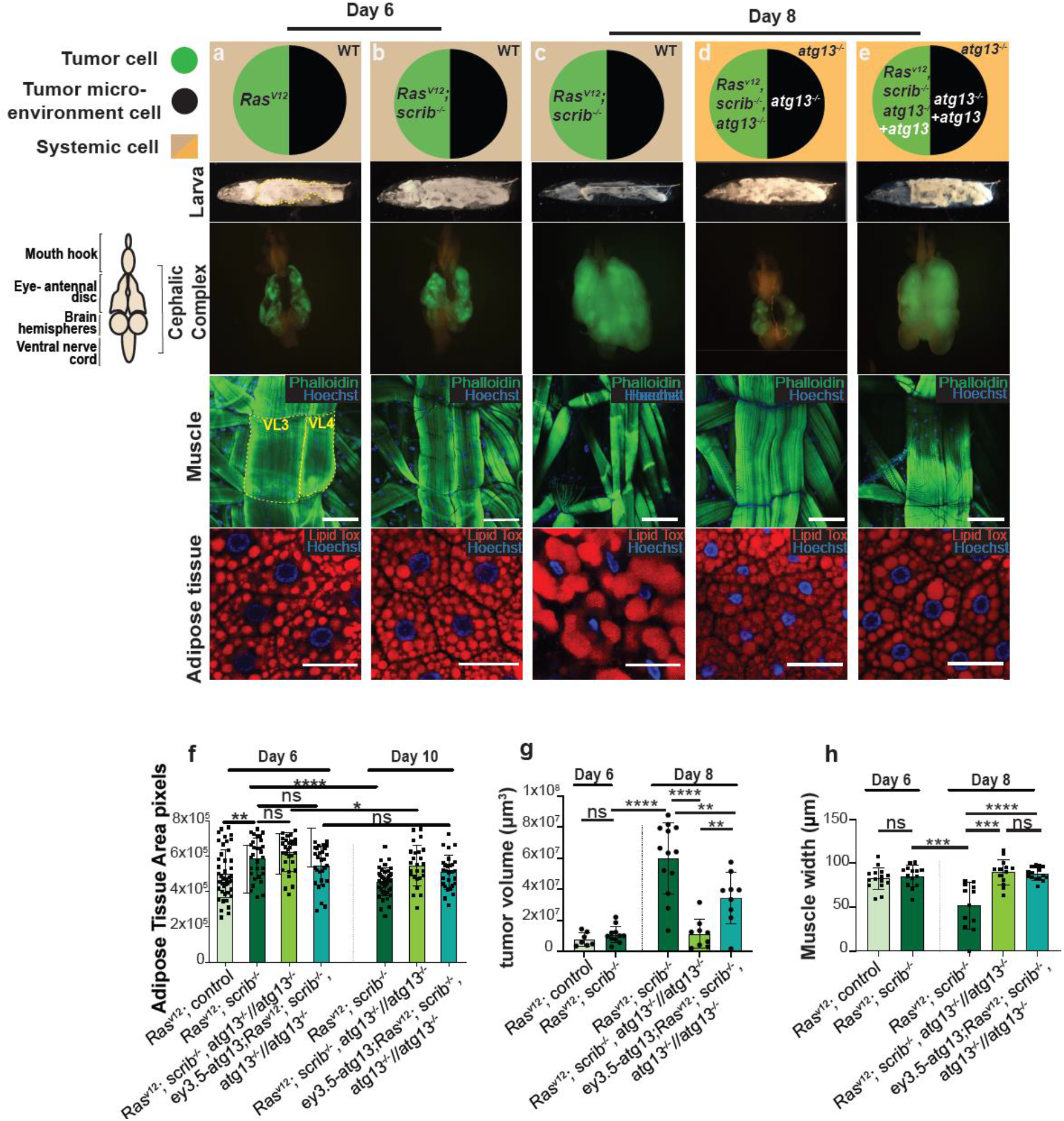
Autophagy is required for systemic wasting. Cartoon (top) illustrates the genotypes of the larvae, eye-antennal disc (EAD, circle: tumor cells in green and microenvironment cell in black), systemic cells are illustrated as a square (wild type cells in light brown and *atg13* mutant cells in light orange) at indicated ages. Cartoon (left) illustrates the structure of cephalic complex attached to mouth hook. Representative images of Larva (image of larva using the backlight of microscope), cephalic complex (green highlights the GFP-labelled tumor clones), muscle (Phalloidin in green stains actin and Hoechst in blue stains nucleus) and adipose tissue (Lipid Tox in red stains for lipid droplets, Hoechst stains nucleus) from top to bottom. (**a**) *Ras^V12^; ctrl* tumor bearing animal at day 6. (**b**) *Ras^V12^; scrib*^-/-^ tumor bearing animal at day 6. (**c**) *Ras^V12^; scrib*^-/-^ tumor bearing animal at day 8. (**d**) *Ras^V12^, scrib^-/-^, atg13^-^7/atg13^-/^* at day 8. (**e**) *Ras^V12^, scrib^-/-^, atg13^-^7/atg13^-/^* animal complemented with transgenic *atg13*, rescuing the tumor growth, at day 8. (**f**) Quantification of area of space occupied with lobes of fat body within larval segments 4 to 8 (shown in yellow dashed line in 2a), of animals bearing tumors at day 6 *Ras^V12^; ctrl* (n=45)*, Ras^V12^; scrib*^-/-^ (n=30), *Ras^V12^, scrib^-/-^, atg13//atg13*^-/-^ (n=30) and *ey3.5- atg13; Ras^V12^, scrib^-/-^, atg13^-/-^//atg13*^-/-^ (n=29) and at day 10 *Ras^V12^; scrib*^-/-^ (n=40), *Ras^V12^, scrib^-/-^, atg13^-^7/atg13^-/^* (n=25) and *ey3.5-atg13; Ras^V12^, scrib^-/-^, atg13^-/-^ //atg13*^-/-^ (n=28). (**g**) Quantification of tumor volumes day 6 *Ras^V12^; ctrl* (n=7)*, Ras^V12^; scrib*^-/-^ (n=11) and day 8 *Ras^V12^; scrib*^-/-^ (n=13), *Ras^V12^, scrib^-/-^, atg13^-^7/atg13^-/^* (n=9) and *ey3.5-atg13; Ras^V12^, scrib^-/-^, atg13^-/-^//atg13*^-/-^ (n=9). (**h**) Quantification of Ventral Longitudinal muscle 4 (VL4) (shown in yellow dashed line in 2a) of third segment of larvae carrying at day 6 *Ras^V12^; ctrl* (n=15)*, Ras^V12^; scrib*^-/-^ (n=14), and at day8 *Ras^V12^; scrib*^-/-^ (n=11), *Ras^V12^, scrib^-/-^, atg13-^/-^//atg13*^-/-^ (n=12) and *ey3.5-atg13; Ras^V12^; scrib^-/-^, atg13^-/-^//atg13^/-^* (n=14). (**i**) Quantification of dry weight excluding tumor of *Ras^V12^; scrib^-/-^, atg13^-/-^//atg13*^-/-^at day 6 (n=8) and day 10 (n=7). (**j**) Quantification of larval motility measured by crawling distance and crawling pattern for *Ras^V12^; scrib^-/-^, atg13^-/-^//atg13*^-/-^ at day 6 (n=15) and at day 8 (n=15), each colored line represents a single larva. Values depict mean±s.e.m. of minimum three independent pooled experiments. ns, not significant, \**P* ≤ 0.05, \*\**P* ≤ 0.01, \*\*\**P* ≤ 0.0001 and \*\*\*\**P* < 0.0001, from unpaired, two-tailed test. Scale bar: muscles 100 μm and adipose tissue 50 μm.

### Systemic autophagy mediates organ wasting

Earlier work showed that larvae carrying *Ras^V12^; scrib*^-/-^ malignant tumors, but not *Ras^V12^* benign tumors display a systemic autophagy stress response in muscle, gut and fat body^4^ To validate these finding we assessed autophagic activity using the autophagy flux reporter mCherry-Atg8a. Increased lysosomal processing of mCherry-Atg8a was observed in host tissues of larvae carrying *Ras^V12^; scrib*^-/-^ tumors from day 6 to 8 (Expanded View Fig. 2a) indicating an escalation in systemic tissue autophagy in this model as tumor growth proceeds. Increased autophagy alongside proteasomal turnover may be responsible for proteolysis and muscle atrophy. We therefore asked whether autophagy is involved in organ wasting. The autophagy initiation complex Atg13/Atg1/FIP200 and Atg14-containing PIK3-C3 lipid kinase complex (Atg14/Vps15/Vps34/Atg6), act sequentially during autophagy initiation. As expected, fully mutant *atg13* animals carrying *Ras^V12^; scrib*^-/-^ tumors, did not display autophagy flux and instead accumulated the autophagy cargo protein Ref(2)P/p62 (Expanded View Fig. 2a). Strikingly, *Ras^V12^; scrib^-/-^, atg13^-/-^//atg13*^-/-^ animals displayed a complete reversal of muscle atrophy, weight and motility loss, as well as fat body alterations (Fig. 3a-d, f-h, Expanded View Fig. 2b,c, Supplementary video 8,9). Genomic rescue of *atg13* (*g-atg13*) in *Ras^V12^; scrib^-/-^, atg13^-/-^//atg13*^-/-^ animals reinstated organ wasting, showing that the failure of organ wasting was indeed due to *atg13* loss (Expanded View Fig. 2d,e). *Ras^V12^; scrib^-/-^, atg14^-/-^//atg14*^-/-^ animals also blocked organ wasting and loss of motility, demonstrating that failure of organ wasting is not limited to the autophagy initiation complex (Expanded View Fig. 2f-g). Importantly, as *Ras^V12^; scrib^-/-^, atg13^-/-^//atg13*^-/-^ also show reduced tumor growth, we rescued tumor growth by local *atg13* expression in the tumor and microenvironment of the eye-antennal disc only. This led to rescue of tumor growth to 60% as previously reported, but failed to cause systemic atrophy, (Fig. 3e, f-h). Thus, systemic autophagy drives organ wasting and tumor growth is dependent upon local as well as systemic autophagy.

### Autophagy mediates nutrient mobilization

Autophagy mediates nutrient recycling upon starvation at the cell level through lysosomal turnover and recycling of lipids, sugars, amino acids and nucleotides that may further be secreted into circulation^1,11,12^. We therefore investigated whether autophagy can mediate systemic nutrient release in this model. The levels of metabolites in the cell free hemolymph (serum) were measured using LC/MS analysis.

Serum levels of amino acids; arginine, proline, glutamine, serine, alanine, lysine as well as the insect serum sugar, trehalose, were significantly increased in animals with *Ras^V12^; scrib*^-/-^ tumors relative to age-matched controls (Fig. 4a). This was apparent already at day 6, before onset of significant and exponential tumor growth and reduced feeding. Conversely, glutamate and methionine were lower (Fig. 4a). By day 8, serum amino acid and trehalose levels increased further, including glutamate and methionine before tapering off at day 10 while glucose peaked (Fig. 4a, Expanded View Fig 3a). Thus, serum nutrient levels increase during organ atrophy suggesting that autophagy may mediate nutrient release into circulation. To test this, we compared *Ras^V12^; scrib*^-/-^ tumor-carrying animals with and without autophagy. Trehalose, arginine, glutamine, proline, tyrosine, serine and asparagine showed significant higher levels in autophagy-competent *Ras^V12^; scrib*^-/-^ animals versus *atg13*-mutant *Ras^V12^; scrib*^-/-^ tumor carrying animals at day 6 (Fig. 4b) and this was followed by the majority of amino acids except tyrosine at day 8 (Expanded View Fig. 3a). This supports the hypothesis that autophagy is required for increased serum nutrient levels in tumor-ridden animals. Indeed, we could not detect significant differences in serum nutrient levels between *Ras^V12^; scrib^-/-^, atg13^-/-^//atg13*^-/-^ and *Ras^✓12^*control animals at day 6 (Expanded View Fig. 3b). In order to infer which metabolites are changed early on during organ wasting in an autophagy-dependent manner, we performed three pairwise comparisons at day 6. Metabolites that increased significantly (FDR < 0.05) upon wasting 1) Ras^V12^;scrib^-/-^ vs Ras^V12^, reversed when autophagy was inhibited 2) Ras^V12^;scrib^-/-^ Atg13^-/-^ vs Ras^V12^;scrib^-/-^ and not altered in autophagy mutants alone 3) Ctrl Atg13^-/-^ vs Ctrl. This set of criteria identified arginine, asparagine, glutamine, proline, and trehalose to respond early and strong at the onset of muscle wasting (Fig. 4c).

**Figure 4.**
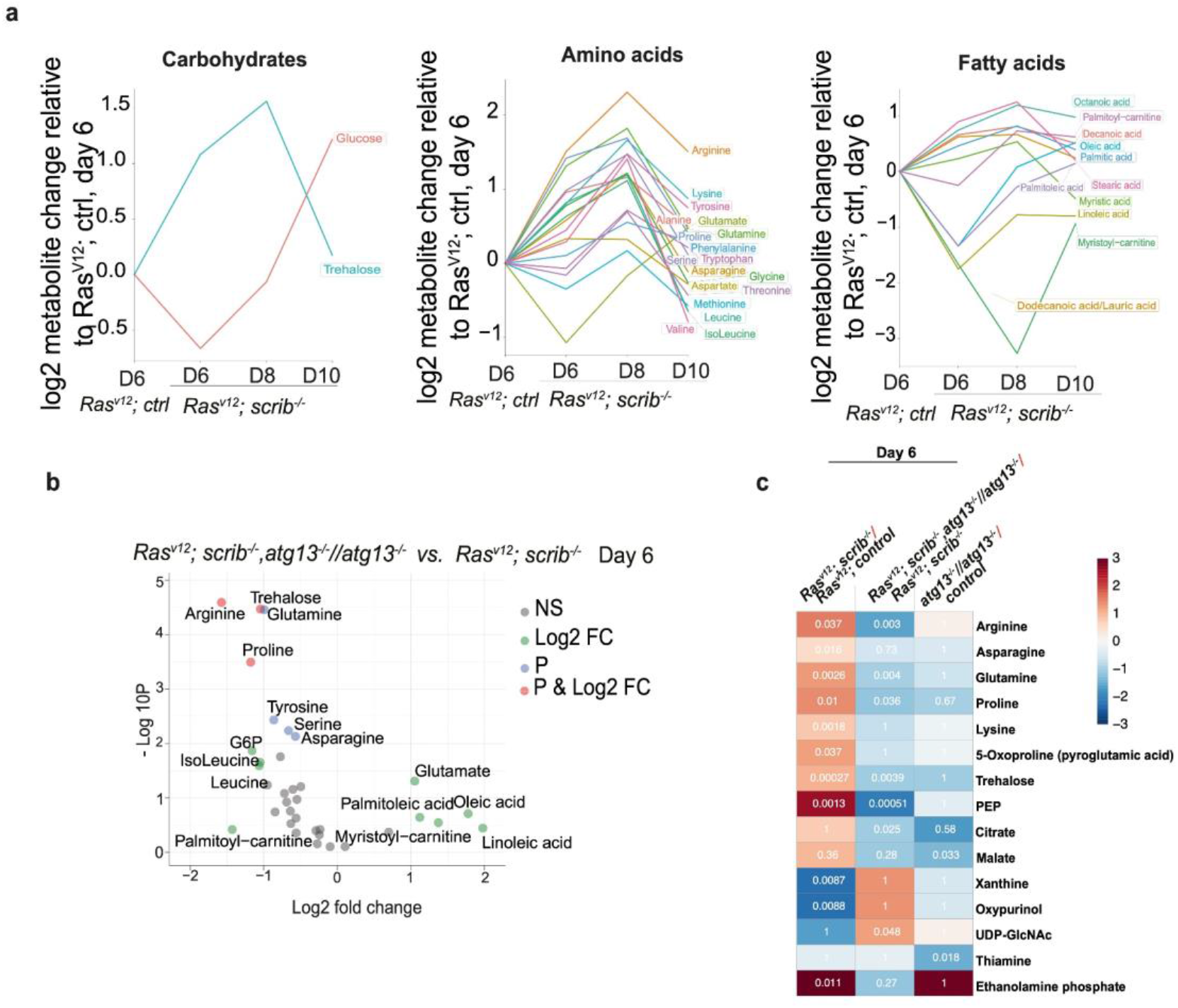
Autophagy-driven wasting releases metabolites into circulation. **(a)** Changes in groups (carbohydrates, amino acids and fatty acids) of storage metabolites measured in circulating haemolymph with progressing wasting, shown as log2 changes measured by LC-MS and calculated per larvae relative to *Ras^V12^; ctrl* at day 6. **(b)** Volcano plot showing autophagy-dependent changes to amounts of circulating metabolites at day 6. X-axis shows the log2 of fold change of *Ras^V12^;atg13 ^-/-^scrib*^-/-^ // *atg13^7-/-^* vs. *Ras^V12^; scrib*^-/-^, y-axis shows −log10 p-value, calculated by t-test. Metabolite names are shown for metabolites with log2(FC) >+ 1 and/or −log10(P) < 2. Green points indicate log2(FC) >+ 1, blue indicate −log10(P) < 2 and red indicates for above both thresholds. **(c)** Autophagy-driven wasting releases metabolites into circulationof 113 reliably detected metabolites, those showing significant differences in any of the three comparisons are shown. Color indicates the log2(fold change) difference and the numbers show the p-value of the comparison. The statistical test to define significance was FDR-adjusted t-test p-value < 0.05

In flies, autophagy cooperates with enzymatic breakdown of muscle glycogen via glycogen phosphorylase in response to starvation, raising the possibility that autophagy may contribute to glycogen breakdown and serum sugar mobilization^13^. Visualizing glycogen by immunofluorescence staining in muscle and fat body, or biochemically measuring total tissue sugar showed that glycogen stores are depleted in day 8 *Ras^V12^, scrib*^-/-^ animals relative to day 6 while serum trehalose levels rise (Fig. 5a-e, Fig. 4c). Both glycogen stores and trehalose levels normalized in *Ras^V12^; scrib^-/-^, atg13^-^7/atg13^-^’* animals (Fig. 5a-e and Expanded View Fig 3c). Taken together, this provides *in vivo* evidence that autophagy mediates somatic organ wasting and mobilization of nutrients in the form of sugars and amino acids and that mobilization initiates before reduced feeding and onset of exponential tumor growth.

**Figure 5.**
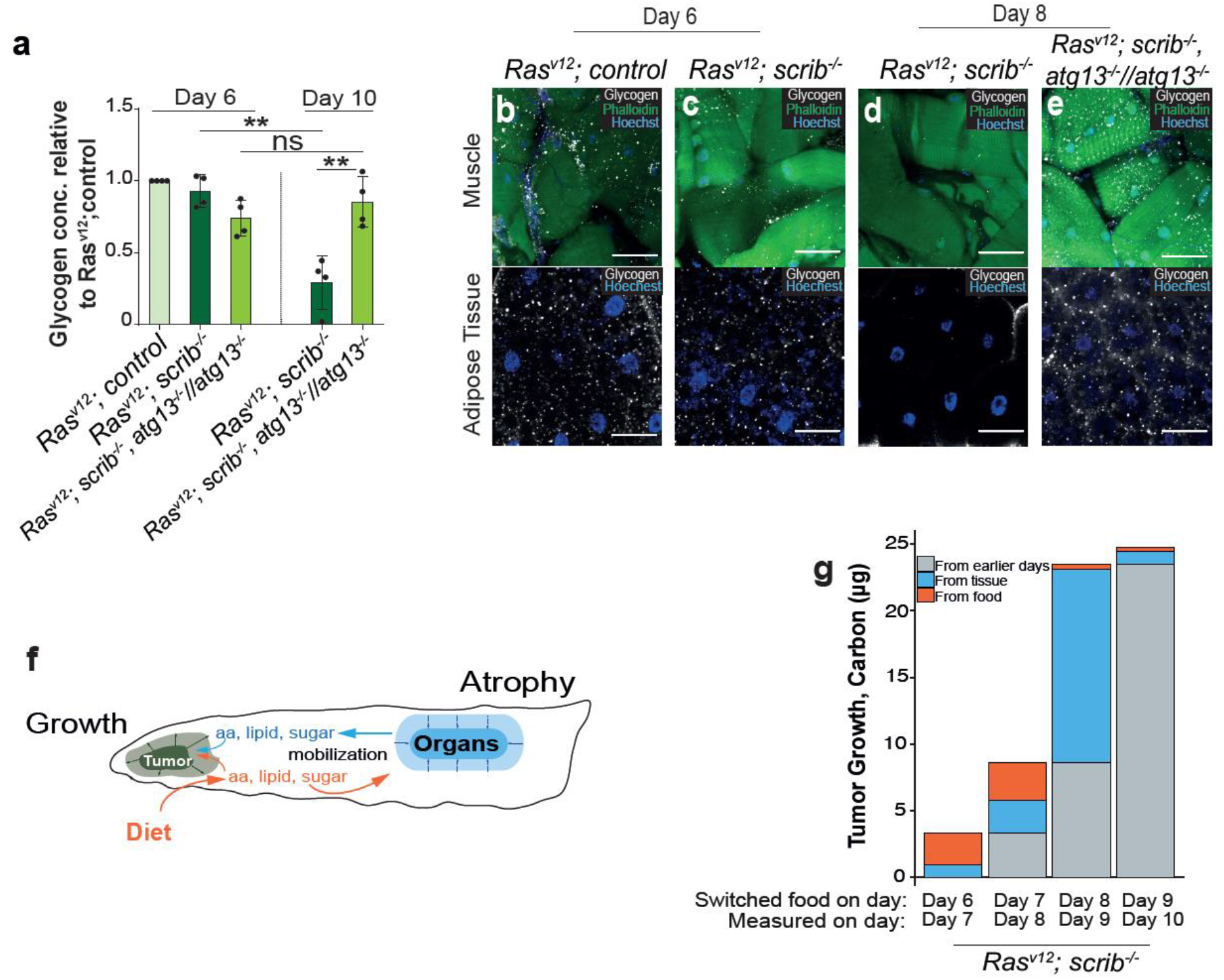
Host-derived nutrients contribute to tumor biomass. **(a)** The amount of glycogen in the whole larvae measured by biochemical assay (n=4), normalized to *Ras^V12^; ctrl* at day 6, and per number of larvae. **(b-e)** Representative confocal images of muscle and adipose tissue of larvae carrying *Ras^V12^; ctrl, Ras^V12^; scrib*^-/-^ and *Ras^V12^; scrib^-/-^, atg13^-/-^//atg13^/-^* showing glycogen levels (white) at day 6 and day 8. (f) Cartoon illustrating tumor growth (green) incorporation of molecules derived from food (in orange) or from host tissues (in blue). **(g)** Sources of carbon incorporated into tumor biomass were differentiated by changing the isotopic carbon content of the food source 25 hours before measuring the total carbon content and isotopic ratio of the tumor. Values depict mean+s.e.m. of minimum three independent pooled experiments. ns, not significant, \**P* ≤ 0.05, \*\**P* ≤ 0.01 and \*\*\*\**P* < 0.0001, from unpaired, two-tailed test. Scale bar: muscles 100 μm and adipose tissue 50 μm.

### Host-derived nutrients constitute the main source of tumor biomass

These findings raised the tantalizing possibility that autophagy mediates nutrient release from host tissues that may further be utilized by the tumor for growth. To test this hypothesis, we sought to differentiate whether biomass of the growing tumor was derived from ingested food or nutrients released from host organ stores. As both sugars and amino acids were mobilized as a group, we chose to follow transfer of carbon using natural abundance Stable Isotope Ratio Mass Spectrometry (SIRMS) using a recently developed protocol^14^. We generated two types of fly food containing different ^13^C/^12^C ratios derived from plants with differences in ^13^C vs ^12^C incorporation from CO_2_ during photosynthesis. C4-type carbon fixation (e.g. corn, sugar cane) incorporates relatively more ^13^C-containing CO_2_ than does C3-type plants (e.g. potato, sugar beet)^14–16^ Larvae raised on C4-based vs C3-based food revealed segregated ^13^C/^12^C values in keeping with the isotopic composition of the food they were raised on (Expanded View Fig 4a). To measure incorporation of carbon from different sources into the growing tumor, we switched the food source 25 hours before measuring the isotopic carbon ratio as well as total carbon mass of the cephalic complex. The measured change in mass and carbon composition during those 25 hours then reveal the relative amount of carbon that has been added to the tumor from ingested food or host organs. We find that early tumor growth sources a majority of biomass carbon from the food, but that carbons are increasingly incorporated from the host tissues as tumor growth progresses (Fig 5f, g, Expanded View Fig 4b).

## Discussion

Earlier studies in adult flies have demonstrated ovary atrophy in adult flies carrying allografted *Ras^V12^; scrib*^-/-^ or *Ras^V12^;dlg*^-/-^ tumors derived from the larval eye disc or in animals with adult stem cell tumors in the gut driven by defective Hippo-Warts signaling. Ovary atrophy is mediated by IMPL2, an insulin-binding peptide produced by the tumor. Muscles showed altered mitochondrial morphology, but muscle wasting was not reported in either model. Detailed μ–CT and volumetric calculations presented here demonstrate that muscle wasting occurs in response to *Ras^V12^; scrib^-/-^*, but that the fat body volume is retained with increased signs of steatosis. Muscle wasting was also reported to be induced by fibroblast growth factor secretion in a *Ras^V12^, Csk-IR* larval model while this paper was being revised^17^ In both models, a concomitant increase in amino acid serum levels is observed raising the question of what is the mechanism for nutrient mobilization.

Autophagy captures intracellular material and delivers it to the lysosome for degradation and recycling of amino acids, sugars, fatty acids and nucleotides to sustain metabolism and enable survival during starvation in mice and flies It is well established that autophagy is required in tumor cells to uphold tumor cell metabolism and mitochondrial functionality cell intrinsically. More recently, non-autonomous effects of autophagy in cancer progression have been revealed. In mouse and *Drosophila, Ras-*driven tumors stimulate neighboring cells of the microenvironment to activate autophagy to support tumor growth^2,4^. Pancreatic stellate cells secrete asparagine and alanine in response to tumor cells and alanine taken up by the tumor was shown to functionally support tumor growth by entering central TCA metabolism^2^. Beyond the microenvironment, systemic depletion of autophagy leads to strong decline in serum arginine due to the release of arginase from the liver that in turn restricts growth of arginine-auxotrophic tumors^19^ Whether autophagy may have a broader effect on physiology and tumor growth when cancer leads to systemic organ wasting is so far unknown. The results herein support a model where autophagy supports tumor growth from peripheral tissues in a fly model^4^. As autophagy mediates massive nutrient release, we favor a model in which autophagy supports tumor growth by bulk nutritional provisioning through amino acid and sugar mobilization. This is supported by measurements showing that biomass increase is mainly derived from host carbon sources in this model. We cannot rule out other autophagy-mediated mechanisms that may also support growth and that the tumor is opportunistically incorporating the released molecules. Given the similarities between human cancer cachexia processes and the findings presented herein, autophagy inhibition may also be an effective way to mitigate symptoms of cachexia and increase quality of life.^18^

---

**Expanded View Figure 1.**
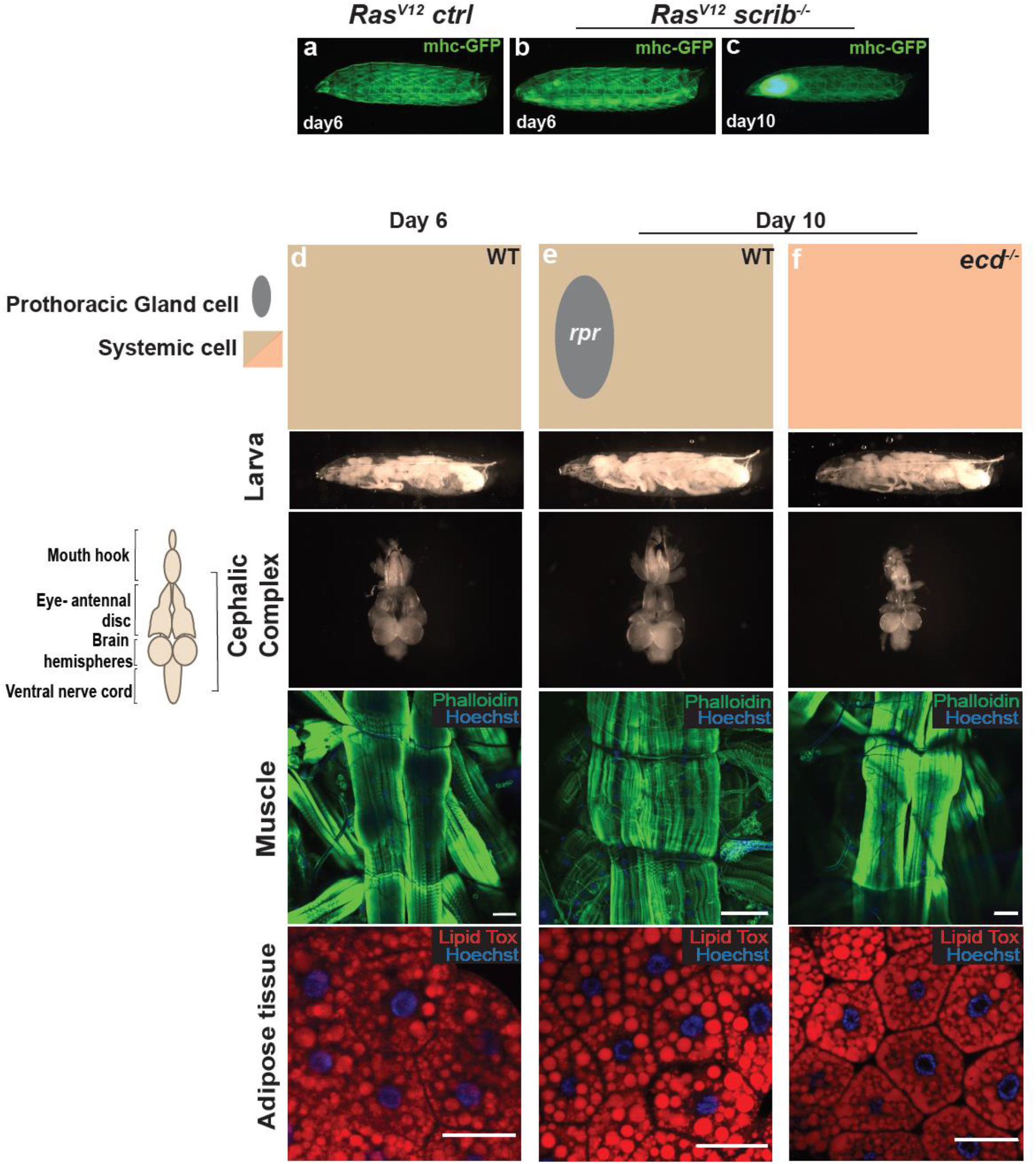
Tumor-induced organ wasting. **(a-c)** Representative images of larvae with GFP-labelled muscles at day 6 and 10. **(d-f)** Cartoon (top) illustrates the genotypes of the larvae, prothoracic gland (grey oval), systemic cells are illustrated as a square (wild type cells in light brown and ecdysone mutant cells in light red) at day 6 and day 10. Cartoon (left) illustrates the structure of cephalic complex attached to mouth hook. Larva (image of larva using the backlight of microscope), cephalic complex (no tumors), muscle (Phalloidin in green stains actin and Hoechst in blue stains muscle nuclei) and adipose tissue (Lipid Tox in red stains for lipid droplets, Hoechst stains cell nuclei) from top to bottom. **(d)** Wild type larva (w^1118^) control, **(e)** spok-Gal4,UAS-Dcr2.D;UAS-rpr larva that linger due to lack of cells expressing ecdysone hormone and **(f)** ecd^1ts^ (ecdysoneless), larva that lingers due to ecdysone deficiency. Scale bar: muscle, 100 μm and adipose tissue, 50μm.

**Expanded View Figure 2.**
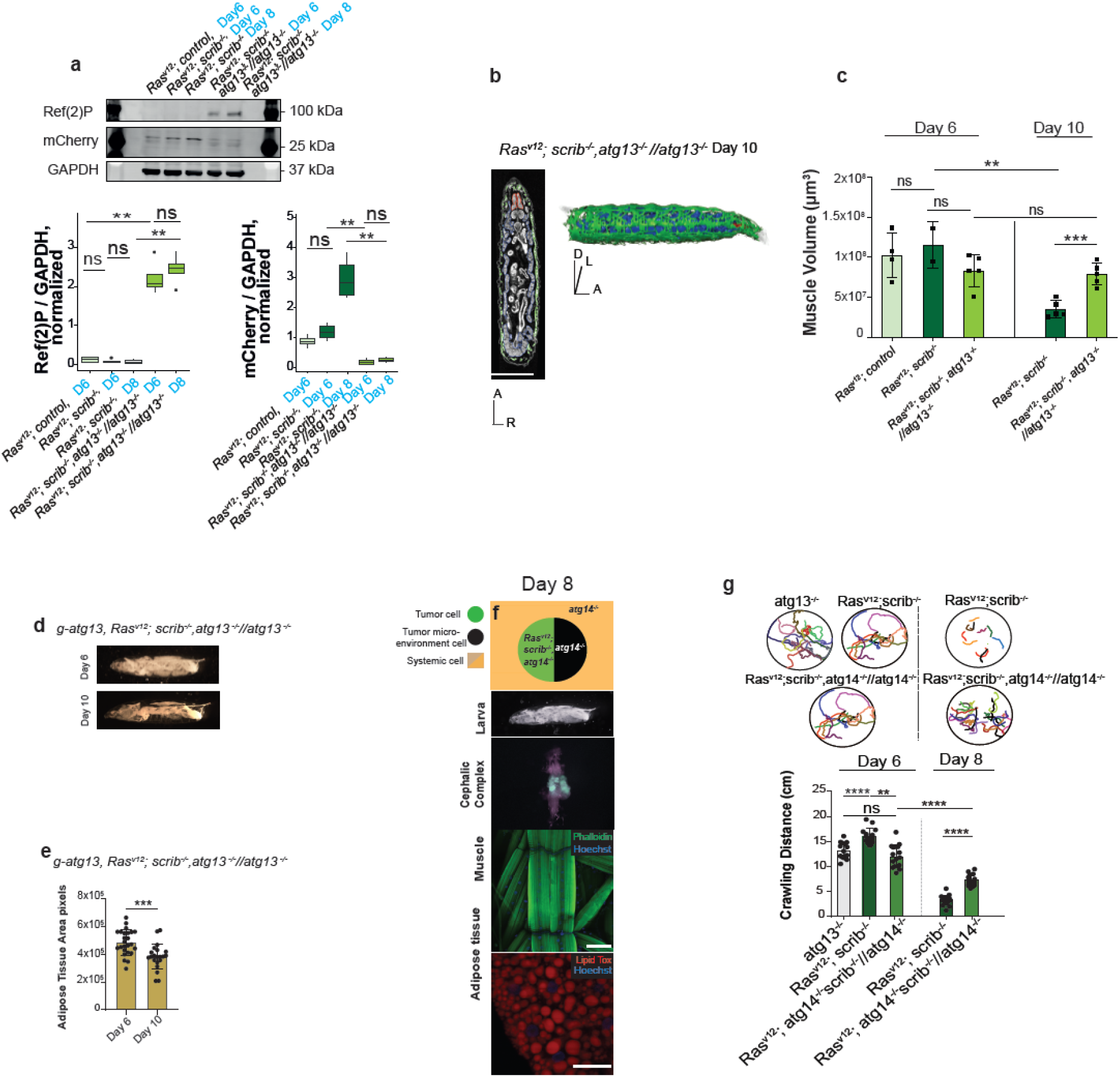
Autophagy is required for systemic wasting. **(a)** Western blot analysis and quantification of Cherry-Atg8a processing and Ref(2)P accumulation in larval body wall muscles, representative of four independent experiments. Quantifications are normalized to the mean of signals for a given band in each independent experiment. **(b)** Representative 2D μ-CT scan and representative 3D rendering of of *Ras^V12^; scrib^-/-^, atg13:^/-^//atg13^-/-^larva* at day 10. Muscle (green), fat body (blue) and eye-antennal discs/tumor (red), Anterior (A), Dorsal (D), Left (L), Right (R). **(c)** Quantifications of muscle volumes larvae carrying at day 6 *Ras^V12^; ctrl* (n=4)*, Ras^V12^; scrib*^-/-^ (n=2), *Ras^V12^; scrib^-/-^, atg13-^/-^//atg13*^-/-^ (n=5) and at day 10 *Ras^V12^; scrib*^-/-^ (n=5), *Ras^V12^; scrib^-/-^, atg13ŕ//atg13^-^’* (n=5). **(d)** Representative images of whole larva at days 6 and 10 of *g-atg13, Ras^V12^; scrib^-/-^, atg13^-/-^//atg13*^-/-^ **(e)** Quantification of space within segments 4-8 occupied by white adipose tissue in a whole/intact larva, shown in pixels **(f)** Cartoon (top) illustrates the genotypes of the larva, eye-antennal disc (EAD, circle: tumor cells in green and microenvironment cell in black), systemic cells are illustrated as a square. Representative images of *Ras^V12^; scrib^-/-^, atg14^-/-^//atg14^-/-^larva* (image of larva using the backlight of microscope), cephalic complex (green highlights the GFP-labelled tumor clones), muscle (Phalloidin in green stains actin and Hoechst in blue stains nucleus) and adipose tissue (Lipid Tox in red stains for lipid droplets, Hoechst stains nucleus) from top to bottom. **(g)** Quantification of larval motility measured by crawling distance and crawling pattern for *atg13*^-/-^ larvae (n=12), *Ras^V12^; scrib^-/-^ (same data that shown in Figure 2.) Ras^V12^; scrib^-/-^, atg14^-/-^//atg14*^-/-^ at day 6 (n=15) and at day 8 (n=16), each colored line represents a single larva. Values depict mean±s.e.m. of minimum three independent pooled experiments, (except for Exp. View Fig 2c, *Ras^V12^; scrib*^-/-^ at day 6) ns, not significant, \**P* ≤ 0.05, \*\**P* ≤ 0.01 and \*\*\**P* ≤ 0.0001 or as indicated in the figure (b), from unpaired, two-tailed test. Scale bar: muscles 100 μm and adipose tissue 50 μm.

**Expanded View Figure 3.**
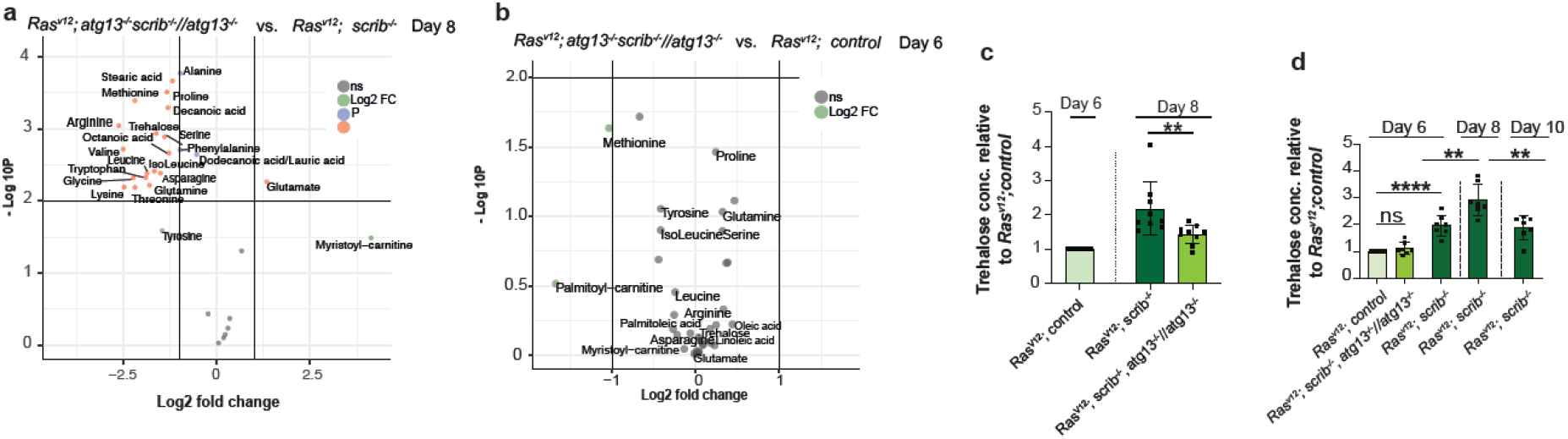
Autophagy-driven wasting releases metabolites into circulation. **(a)** Volcano plot showing autophagy-dependent changes to amounts of circulating metabolites in wasting larvae at age 8. X-axis shows the log2 of fold change of *Ras^V12^; scrib^-/-^, atg13^-/-^//atg13*^-/-^ vs. *Ras^V12^; scrib*^-/-^, y-axis shows −log10 p-value, calculated by t-test. Metabolite names are shown for metabolites with log2(FC) >+ 1 and/or −log10(P) < 2. Green points indicate log2(FC) >+ 1, blue indicate - log10(P) < 2 and red indicates for above both thresholds. **(b)** Volcano plot showing the amounts of circulating metabolites in *Ras^V12^; scrib^-/-^, atg13^-/-^//atg13*^-/-^ vs. *Ras^V12^; ctrl* at day 6. **(c)** The amount of trehalose measured by biochemical assay in whole larvae (n=9) in *Ras^V12^; scrib*^-/-^ and *Ras^V12^; scrib^-/-^, atg13^-/-^//atg13*^-/-^ at day 8, normalized to *Ras^V12^; ctrl* at day 6, and per number of larvae. (d) The amount of trehalose measured by biochemical assay of the whole larvae (n=7), in *Ras^V12^; scrib^-/-^, atg13^-/-^//atg13*^-/-^ at day 6 and *Ras^V12^; scrib*^-/-^ at day 6, 8 and 10 normalized to *Ras^V12^; ctrl* at day 6, and per number of larvae. Values depict mean±s.e.m. of minimum three independent pooled experiments; ns, not significant, \*\**P* ≤ 0.01\*\**P* ≤ 0.01and \*\*\*\**P* < 0.0001, from unpaired, two-tailed test.

**Expanded View Figure 4.**
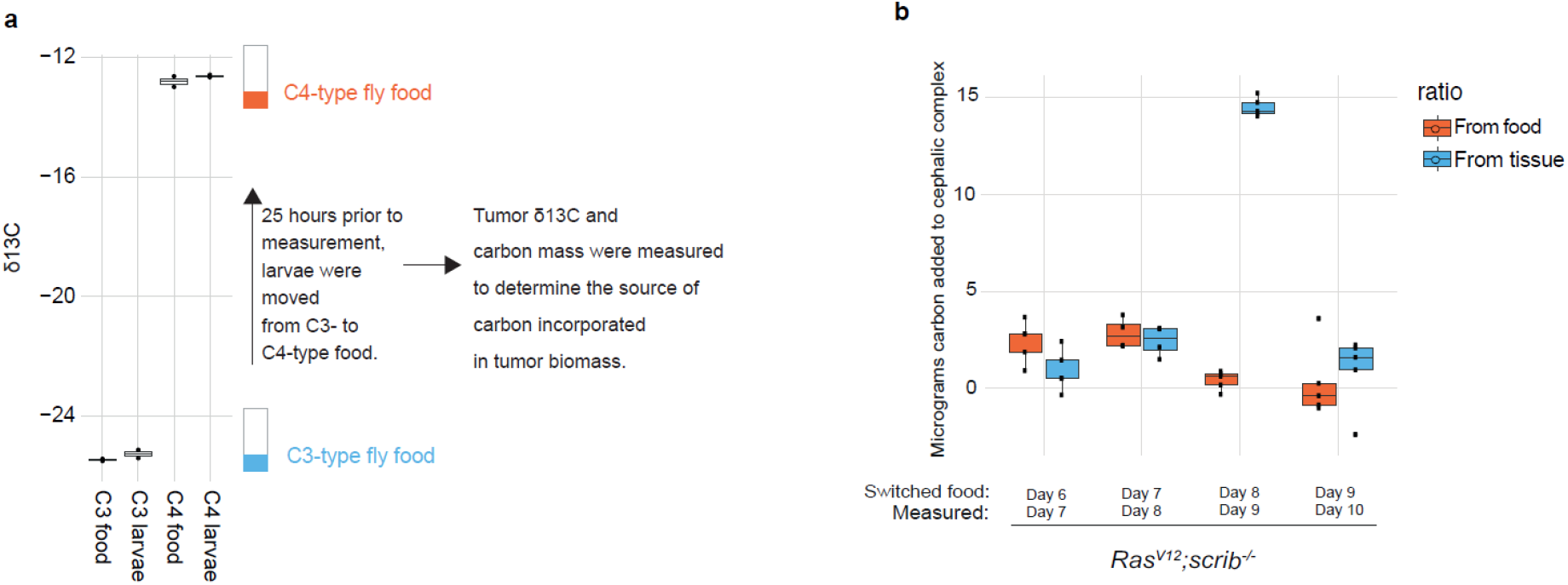
Tumor incorporation of carbon isotopes. Determining the source of carbon used for tumor biomass expansion. (a) δ13C (representing 13C/12C) measurements of two different food types, either with C3- or C4-type plant-derived nutrients. Measurements of whole larvae growing on these food types for are also shown along with a brief description of the methodology to determine the source of carbon used for tumor growth. Two measurements are shown for all samples. (b) Carbon incorporated into the cephalic complex biomass from food or host tissues were determined by moving RasV12; scrib-/- larvae at indicated stages tumor development from C3- to C4-type food and then measuring the carbon composition as well as carbon mass 25 hours later.

## Supplementary files

### VIDEOS

**Supplementary Video 1.** 2D μ-CT scan of a *Ras^V12^; ctrl* larva at Day 6, showing the entire range of projection images acquired from two different perspectives. Muscle (green), fat body (blue) and eye-antennal discs/tumor (red) are outlined. *Top:* Movie begins at the right side of the larva and moves towards the left side. *Bottom:* Movie begins at the dorsal surface and moves towards the ventral surface (top to bottom). Scale Bar: 1mm; Anterior (A), Dorsal (D), Right (R).

**Supplementary Video 2.** 2D μ-CT scan of a *Ras^V12^; scrib*^-/-^ larva at Day 6, showing the entire range of projection images acquired from two different perspectives. Muscle (green), fat body (blue) and eye-antennal discs/tumor (red) are outlined. *Top:* Movie begins at the right side of the larva and moves towards the left side. *Bottom:* Movie begins at the dorsal surface and

**Supplementary Video 3.** 2D μ-CT scan of a *Ras^V12^; scrib*^-/-^ larva at Day 10, showing the entire range of projection images acquired from two different perspectives. Muscle (green), fat body (blue) and eye-antennal discs/tumor (red) are outlined. *Top:* Movie begins at the right side of the larva and moves towards the left side. *Bottom:* Movie begins at the dorsal surface and moves towards the ventral surface (top to bottom). Scale Bar: 1mm; Anterior (A), Dorsal (D), Left (L).

**Supplementary Video 4.** 3D rendering of *Ras^V12^; ctrl* larva at Day 6 showing muscle (green), fat body (blue) and eye disc/tumor (red). Movie begins at the right side of the larva and clips towards the left side before reversing direction. Movies towards the ventral surface (top to bottom). Scale Bar: 1mm; Anterior (A), Dorsal (D), Right (R).

**Supplementary Video 5.** 3D rendering of *Ras^V12^; scrib*^-/-^ larva at Day 6 showing muscle (green), fat body (blue) and eye disc/tumor (red). Movie begins at the right side of the larva and clips towards the left side before reversing direction.

**Supplementary Video 6**. 3D rendering of *Ras^V12^; scrib*^-/-^ larva at Day 10 showing muscle (green), fat body (blue) and eye disc (red). Movie begins at the right side of the larva and clips towards the left side before reversing direction.

**Supplementary Video 7.** 3D rendering of *Ras^V12^; scrib*^-/-^ larva tumor (red) and central nerve system (yellow). Movie shows the growth of tumor in time, through day 6 to day 10.

**Supplementary Video 8.** 2D μ-CT scan of a *Ras^V12^; scrib*^-/-^ larva at Day 10 (top) and *Ras^V12^; scrib^-/-^, atg13^-/-^//atg13*^-/-^ larva at Day 10 (bottom), showing the entire range of projection images acquired. Muscle (green), fat body (blue) and eye-antennal discs/tumor (red) are outlined. Movie begins at the dorsal surface and moves towards the ventral surface (top to bottom). Scale Bar: 1mm; Anterior (A), Right (R).

**Supplementary Video 9.** 3D rendering of *Ras^V12^; scrib*^-/-^ larva at Day 10 (top) and *Ras^V12^; scrib^-/-^, atg13^-/-^//atg13*^-/-^ larva at Day 10 (bottom) showing muscle (green), fat body (blue) and eye disc/tumor (red). Movie begins at the right side of the larva and clips towards the left side before reversing direction. Anterior (A), Dorsal (D), Left (L).

## Materials and methods

### Fly Husbandry and fly food

Stocks and crosses were kept at 25 °C on standard potato mash fly food containing 32.7 g dried potato powder, 60 g sucrose, 27.3 g dry yeast, 7.3 g agar, 4.55 ml propionic acid, and 2 g nipagin per liter, resulting in a final concentration of 15.3 g l^-1^ protein and 6 g l^-1^ sugar, unless otherwise indicated. Additional yeast paste was added to the vials.

Temperature sensitive crosses (Figure 4c-h), were set up in 18°C on standard fly food. Third instar larval stage L2 larvae were collected and transferred to 30 °C incubator for induction of selected genes, under the temperature sensitive promoter.

### Immunofluorescence staining and microscopy

Larval cephalic complexes, muscles and fat bodies were dissected in PBS and fixed in PBS containing 4% paraformaldehyde for 30 min at room temperature. Samples were washed twice with PBS and blocked with PBT (PBS containing 0.1% Triton X-100 and 3% BSA). Primary antibody was incubated over night at +4°C, followed by two washes with and secondary antibodies, incubated at room temperature for 1h. Tissue samples were washed and mounted in Vectashield mounting medium (H-1000) and observed with a Zeiss LSM 710 or LSM 780 confocal microscope. Hoechst 33342 (Life technologies, H3570, final concentration 5 μg ml^-1^) and Alexa Fluor-488/647 Phalloidin (H3572, 1:200) were added to stain nuclei and actin, respectively and Lipid Tox Red (Thermo Fisher Scientific, 1:1000). Antibodies used in this study: antiglycogen monoclonal glycogen antibody (1:100, mouse, IgM, Generous gift from Otto Baba).

### Metabolite analysis

20 larvae were washed thoroughly with cold Saline (0.9% NaCl solution), ensuring no traces of food was left on the larval body wall. Larvae were dried completely of saline solution by placing them on Kim Tech paper wipes. Larvae were then transferred into pre-cooled 0.5 ml microcentrifuge tube with a 3-4 mm cut at the bottom center vertically, one by one and using forceps tip a hole was made into the head region of the larva, avoiding puncturing the tumor or the gut. The 0.5 ml microcentrifuge tube was then transferred into a pre-cooled 1.5 ml microcentrifuge tube and centrifuged at 1000 x g for 1 min at 4°C for hemolymph extraction. Tube containing the larval carcasses was removed and extracted hemolymph solution was re-centrifuged at *1000 x g, for 5 min* at 4°C for the removal of hemocytes. 5 μl of serum was pipetted into a new pre-cooled microcentrifuge tube avoiding the hemocytes in the bottom, followed by immediate addition of 70μl of the extraction solvent (LC-MS methanol 75 %: LC-MS acetonitrile 25%, Merck, containing labelled standards, Metabolomic QC Kit for Untargeted/Targeted Mass Spectrometry, A Cambridge Isotope Laboratories Company). The solution was mixed well for 5 min. at 4°C. Sample solutions were centrifuged at 14 000 rpm, for 10 min at 4°C. The supernatant was moved into a new Eppendorf tube and kept at −80°C until analysis. LC-MS metabolomics analysis was performed as_described previously^20^ Briefly, Thermo Ultimate 3000 high-performance liquid chromatography (HPLC) system coupled to Q-Exactive Orbitrap Mass Spectrometer (Thermo Fisher Scientific) was used with a resolution of 35,000 at 200 mass/charge ratio (*m*/*z*), electrospray ionization, and polarity switching mode to enable both positive and negative ions across a mass range of 67 to 1000 *m*/*z*. HPLC setup consisted ZIC-pHILIC column (SeQuant; 150 mm x 2.1 mm, 5 μm; Merck), with a ZIC-pHILIC guard column (SeQuant; 20 mm x 2.1 mm). 5 ul of Biological extracts were injected and the compounds were separated with mobile phase gradient of 15 min, starting at 20% aqueous (20 mM ammonium carbonate adjusted to pH 9.2 with 0.1% of 25% ammonium hydroxide) and 80% organic (acetonitrile) and terminated with 20% acetonitrile. Flow rate and column temperature were maintained at 0.2 ml/min and 45°C, respectively, for a total run time of 27 min. All metabolites were detected using mass accuracy below 5 ppm. Thermo Xcalibur was used for data acquisition

TraceFinder 4.1 was used for analysis. Peak areas of metabolites were determined by using the exact mass of the singly charged ions. The retention time of metabolites was predetermined on the pHILIC column by analyzing an in-house mass spectrometry metabolite library consisting of commercially available standards. For data normalization, raw data files were processed with Compound Discoverer 3.0 to obtain total compounds peak area for each sample. Each metabolite peak area value analyzed in the sample was normalized to total measurable ions in the sample.

For further calculations, the ion counts measured by the LC-MS were imported to R (R Core Team, 2019). From the concentration/metabolites peak area that were measured on the LC-MS instrument, the normalized total amount of each metabolite in circulation per larvae was calculated by multiplying by the hemolymph volume of the larvae with the peak area and dividing by the total measurable ions for that sample. Peak area, volumes and total measured ions for all samples are supplied in Supplementary Data 1. To allow log2 (fold change) measurements even if there are metabolites that are not detected in a sample set, all normalized total amounts were added 0.0001 for the log2(fold change) calculations. P values were calculated by the t.test function in R and volcano plots made by the Enhanced Volcano (Kevin Blighe, 2019) library in R

### Trehalose measurement

Three larvae of the indicated ages were collected and washed thoroughly in cold PBS three times. Larvae were then dried on a Kim Tech paper, and transferred into a 1.5 ml microcentrifuge tube containing ice cold lysis buffer 20 μl/larva (Pierce Luciferase Cell Lysis Buffer (2X), Thermo Scientific) and immediately homogenized using a pellet pestle. Samples were then heated for 10 min at 70°C, followed by centrifugation at 14 000 rpm, for 10 min at 4°C. The supernatant was transferred into a new 1.5 ml microcentrifuge tube.

Trehalose amount was measured using Trehalose Assay Kit (Product code: K-TREH, Megazyme Ltd.) according to the manufacturer’s instructions, with the exception of using 20μl of larval sample extraction per 200μl of total sample volume, in 96 well plate. Two aliquots were measured, with and without trehalase enzyme, to measure background glucose amount and trehalose converted to glucose amount. Samples were normalized to the amount of larvae and to the control sample (*Ras^V12^;control*).

### Glycogen measurement

Three larvae of the indicated ages were collected and washed thoroughly in cold PBS three times. Larvae were then dried on a Kim Tech paper, and transferred into a 1.5 ml microcentrifuge tube containing ice cold lysis buffer 30 μl/larva (Pierce Luciferase Cell Lysis Buffer (2X), Thermo Scientific) and immediately homogenized using a pellet pestle. Samples were then heated for 10 min at 70°C, followed by centrifugation at 14 000 rpm, for 10 min at 4°C. The supernatant was transferred into a new 1.5 ml microcentrifuge tube.

Glycogen amount was measured using Glycogen Colorimetric Assay Kit II (Catalog *#:* K648, BioVision, Inc), according to the manufacturer’s instructions, with the exception of using 10μl of larval extract solution in the total volume of 150 μl. An addition sample aliquot was measured without Hydrolysis Enzyme Mix to measure the background glucose level, which was subtracted from the total glucose level (background glucose + glycogen converted to glucose) to obtain glycogen only amount. Samples were normalized to the amount of larvae and to the control sample (*Ras^V12^;control*).

### mCherry flux assay, western blot

Body wall perparations were made from three dissected larvae, added to 30 μl RIPA buffer (150 mM NaCl, 50 mM Tris-HCl pH 7.5, 1% Triton X-100, 1% sodium deoxycholate, 0.2% SDS) with complete protease inhibitor cocktail (Sigma, 11697498001). Lysates were sonicated 10 cycles of 30 seconds at high intensity then debris sedimented by centrifugation 10 minutes, 18000 g at 4 °c. The cleared lysates were normalized to protein content by the Pierce BCA assay kit (Thermo Scientific, 23227) and 8 μg of protein was added from each sample to a 26-well Criterion TGX 4-20% precast gel (Bio-rad, 5671095). Proteins were transferred to an Immun-Blot low flouresence PVDF membrane (Bio-rad, 1620264) using a semi-dry transfer system. Membranes were first blocked in 5% FA-free BSA for 1 hour (Sigma, A7030) then incubated with primary antibodies goat anti-mCherry (Acris, AB0040-200) at 1:1000, mouse anti-GAPDH (Abcam, ab9484) at 1:2500, rabbit anti-Ref(2)p (generated in-house). Membranes were then washed and incubated with fluorescent secondary antibodies for the appropriate species (Li-Cor) using either the 680 nm or 800 nm channels. Finally, membranes were washed and flourescence detected using an Odyssey imaging system (Li-Cor). All membrane incubations and washes were performed in PBS with 0.05% Tween-20 (Sigma, P1379). Membrane band quantification was performed with ImageJ, first marking the band area by the rectangle tool, creating a closed area that selects the band area while excluding any surrounding noise by a straight line then selecting the rectangle by the wand tool which gives the quantified band intensity.

### Larval Locomotor Assays

Larval locomotor assay was conducted to measure distance travelled in the set time of 4 minutes. Individual larva was carefully set on a 15 cm Petri dish containing harden 2% agarose. Larval movement was tracked by drawing over the petri dish cap. Larval travelled distance was measured using fiji and each line was normalized to the cap diameter.

### Larval Dry Weight

Glass microscopy slides were weighed on a ME54 Mettler Toledo scale. 10 larvae were collected at different time points, washed 3x in water to remove any food particles, dried off on Kim wipes and dissected directly on the glass slide to remove the entire transformed brain complex and eye discs. Slides were allowed to dry at room temperature for 24h and weighed again to determine the dry weight of the animals.

### Heat Fixation and imaging

Larva was collected and washed thoroughly in PBS, and transferred into a drop pf 100% glycerol on a microscope slide. To heat fix the larva the slide was placed onto a heat block at 70°C for approximately 10 sec. or until larva immobile. Overview of larval fat body and muscles (FGP-labelled myosin heavy chain (Mhc)) were obtained on a Leica MZ FLIII fluorescence stereomicroscope with a Leica DFC420 camera, using the backlight or fluorescence filters. Dissected cephalic complexes were imaged immediately in a drop of PBS similarly. For the segmentation of fat body occupied space, images were analyzed with Fiji image processing software. Background was subtracted with a rolling ball (radius 400), and images smoothened with Gaussian blur filter (sigma =2). Thresholding was then performed using the “Minimum dark” method.

### Carbon Transfer measured by Stable Isotope Ratios (CATSIR)

C3-based food was prepared with 32.7 g/L potato mash, 60 g/L sucrose from beets and 27.3 g/L dry yeast (*δ^13^C* measured to be similar to C3-type sugars). C4-based food was prepared with 32.7 g/L corn flour, 62 g/L sucrose from sugar cane and 26.3 g/L dry yeast that was expanded on sugar cane sucrose. The sugar cane and yeast content of the C4 food was adjusted to account for the slightly higher protein content and lower carbohydrate content of the corn flour compared to the potato mash, giving a similar final fat, protein and carbohydrate content of the two food variants. Both foods were also added 4.55 ml/L propionic acid (Sigma, P5561), 2 g/L nipagin (Sigma, H5501) and 7.3 g/L agar (AS Pals, 77000). When moving larvae from C3 to C4 food, holes were poked in surface of the new food to give easy access and the larvae were then left for 24+1 hours, assuming 1 hour for the previous C3 food to be absorbed or passed through the gut (Wong et al, Nature Methods, 2008). Before dissection larvae were washed 3 times in water and cephalic complex was dissected in a drop of ultrapure water. The cephalic complex from a single larvae was added to a tin cup (Elemental microanalysis, D1006) and samples were left to dry in a dessicator.

The δ^13^C value and total carbon mass of each sample was determined in triplicate using a Delta V Advantage Isotope Ratio Mass Spectrometer (Thermo Fisher, Bremen, Germany) configured to a Thermo Finnigan 1112 Series Flash Elemental Analyzer at the University of Oslo, Norway. Raw data is reported in standard δ-notation (in units of permil, ‰) and normalized to the Vienna Pee Dee Belemnite (VPDB) scale using three internal lab reference materials (JRICE: δ^13^C = −27.44‰; JHIST: δ^13^C = −8.15‰; and JGLY: δ^13^C = −43.51‰). For all samples, the median standard deviation of the replicate capsules was 0.10‰.

To determine how much carbon is derived from the food or host tissues after moving the larvae from one food to the other, δ^13^C measurement of any given day were compared to a set of measurements of larval cephalic complexes growing on only C3 or C4 food. This comparison is required because the δ^13^C of the cephalic complex with a growing tumor gradually changes as the tumor grows, as is described in more detail in the method publication^14^ First, the relative ratio of carbon derived from food and host was determined by comparing the measured δ^13^C to the C3- and C4-only samples. Then the measured total carbon mass of the cephalic complex was multiplied by this relative ratio to determine the total carbon added from the two different carbon sources.

### μ-CT

#### Labeling

Third instar larvae were collected and transferred to a 1.5ml Eppendorf tube containing 1 ml of 0.5% phosphate buffered saline + Triton-X 100 (0.5% PBST) and incubated for 5 minutes at room temperature. Larvae were then fixed in 1 ml Bouin’s solution (5% acetic acid, 9% formaldehyde, 0.9% picric acid; Sigma Aldrich, St. Lous, MO) for 2 hrs at room temperature. A microdissection needle was then used to poke a small hole in the cuticle at both the anterior and posterior ends of each larva, carefully avoiding any underlying soft tissue. This allows for better penetration of the Bouin’s solution and even fixation of all larval tissues. Fresh Bouin’s solution was then added and larvae were left to incubate another 16-22 hrs at room temperature. Larvae were then washed 3×30 minutes in 1 ml of μ-CT Wash Buffer (0.1M Na_2_HPO_4_, 1.8% Sucrose) and stained with 1 ml of a 0.1N solution of iodine-iodide (I_2_KI, Lugol’s solution) for 1-2 days at room temperature. Larvae were then washed in two changes of ultrapure water and stored at room temperature.

#### Scanning

Individual larva were mounted for scanning using a P10 pipet tip filled with water, wedged inside a small piece of plastic capillary tube that fits tightly in a custom-made brass holder. A dulled 20-gauge needle was used to gently push the larva down in the pipet tip until it became wedged along the wall, which was necessary to prevent sample movement during scanning. Larva were scanned using a SkyScan 1172 desktop scanner controlled by Skyscan software (Bruker) operating on a Dell workstation computer (Intel Xeon X5690 processor @ 3.47GHz (12 CPUs), 50 GB RAM and an NVIDIA Quadro 5000 (4GB available graphics memory) GPU). X-Ray source voltage & current settings: 40 kilovolts (kV), 110 microamps (μA), 4 watts (W) of power. A Hamamatsu 10Mp camera with a 11.54 μm pixel size coupled to a scintillator was used to convert X-rays to photons. Medium camera settings at an image pixel size of 4.5 μm were used for fast scans (~20 minutes), which consists of about 300 projection images that are acquired. Small camera settings at an image pixel size of 2.5 μm and utilizing 360 degrees of sample rotation (~1500 projection images) were used for the slow overnight (O/N) scans. Random movement was set to 10 and frame averaging ranged from 4-8. At these settings, the scanner has a measured resolution of 5.5 μm under optimal imaging conditions (Morales et al., 2016). Medium camera settings (fast scans) provide enough resolution for accurate morphometric analysis of tissues, whereas small camera settings (slow O/N scans) provide high resolution for assessment of tissue/tumor morphology.

#### Reconstruction

Tomograms were generated using NRecon software (Bruker MicroCT, v1.7.0.4). Reconstruction was performed using a built-in shit correction algorithm that uses reference scans to correct for sample drift in the projection images, followed by an iterative application of misalignment and ring artifact reduction algorithms to generate the highest quality images possible (Salmon et al., 2009).

#### Image Analysis & Statistics

Tomograms were visualized and analyzed using Dragonfly (v4.1, Object Research Systems (ORS) Inc, Montreal, Canada, 2019; software available at http://www.theobjects.com/dragonfly) operating on a Dell workstation (Intel Xeon CPU E5-2623 @ 3.00GHz (16 CPUs), 32 GB RAM, NVIDIA Quadro M2000 GPU (20 GB available graphics memory) and Dell Precision T7600 workstation (Intel^®^ Xeon^®^ CPU E5-2620 @ 2.00GHz, 32GB RAM, 64-bit Operating System, 3GB NVIDIA Quadro K4000 GPU). Segmentation of anatomical structures into individual regions of interest (ROIs) was performed manually using thresholded images with the 3D paintbrush function. To eliminate human bias during segmentation, a single threshold value was selected that encompassed the entire structure of interest in all images that were to be compared. This threshold value was then applied to all images and used for segmentation of tissues into ROIs. Small adjustments to this threshold value for some individuals were made when necessary to fully encompass a given tissue to be measured, to account for slight differences in contrast (as a result of X-Ray beam fluctuations) between samples. ROIs were then converted to meshes for quantitative analysis and visualization. The built-in 2D and 3D movie maker was used to generate all movies.

## Acknowledgements

R.K, F.O.F, A.J., C.D. and T.E.R were supported by grants #262652 and #276070 from the Norwegian Research Council (to TER). P.H. S.T. were supported by HSØ grant #40041 (to T.E.R). NMR and TAS are supported by the Division of Intramural Research at the National Institutes of Health/National Heart, Lung, and Blood Institute (1ZIAHL006126 to NMR and 1K22HL137902 to TAS). We thank Anne Simonsen, Harald Stenmark and Jorrit Enserink for critically reading the manuscript. We thank the Laura and Isaac Perlmutter Fund for supporting the Technion’s metabolomics facility, and the Simon Fougner Hartmann’s fund for research support (to T.E.R). This work was supported by Office of Science, Research Council of Norway through its Centers of Excellence funding scheme #223272.

## Author Contributions

R.K, P.H, E.G., W.M.H., A.H.J. and T.E.R. designed the research; R.K., P.H., T.S., I.A., S.T., C.D., A.J., T.A.S, F.O.F., S.W.S., N.S.K, and M.M.R. performed experiments and analyzed the data. E.M.Q. and R.N. performed analysis of μ-CT data. A.B., H.J. and N.M.R. contributed to the conception of the work. R.K., P.H. and T.E.R wrote the manuscript. T.E.R is the leading principal investigator who conceived the project, supervised research and edited the paper.

The authors declare that they have now conflict of interest

## Detailed genotypes

**Figure 1.**

**1a-c:** *y,w,ey-FLP/y,w;UAS-Ras^V12^/act>y^+^>GAL4, UAS-GFP; Frt82B/Frt82B, tub-GAL80 y,w,ey-FLP/y,w;UAS-Ras^V12^/act>y^+^>GAL4, UAS-GFP; Frt82B, scrib^1^/Frt82B, tub-GAL80*

**Figure 2.**

**1a-h:** *y,w,ey-FLP/y,w;UAS-Ras^V12^/act>y^+^>GAL4, UAS-GFP; Frt82B/Frt82B, tub-GAL80 y,w,ey-FLP/y,w;UAS-Ras^V12^/act>y^+^>GAL4, UAS-GFP; Frt82B, scrib^1^/Frt82B, tub-GAL80*

**Figure 3.**

**3a:** *y,w,ey-FLP/y,w;UAS-Ras^V12^/act>y^+^>GAL4, UAS-GFP; Frt82B/Frt82B, tub-GAL80*

**3b,c:** *y,w,ey-FLP/y,w;UAS-Ras^V12^/act>y^+^>GAL4, UAS-GFP; Frt82B, scrib^1^/Frt82B, tub-GAL80*

**3d:** *y,w,ey-FLP/y,w;UAS-Ras^V12^/act>y^+^>GAL4, UAS-GFP; Frt82B, atg13^Δ81^scrib^1^/Frt82B, tub-GAL80, atg13^Δ81^*

**3e:** *y,w,ey-FLP/ey3.5-Hsp70-DmAtg13-GFP;UAS-Ras^V12^/act>y^+^>GAL4, UAS-GFP; Frt82B, atg13^Δ81^scrib^1^/Frt82B, tub-GAL80, atg13^Δ81^*

**3f,g,h:** *y,w,ey-FLP/y,w;UAS-Ras^V12^/act>y^+^>GAL4, UAS-GFP; Frt82B/Frt82B, tub-GAL80*

*y,w,ey-FLP/y,w;UAS-Ras^V12^/act>y^+^>GAL4, UAS-GFP; Frt82B, scrib^1^/Frt82B, tub-GAL80*

*y,w,ey-FLP/y,w;UAS-Ras^V12^/act>y^+^>GAL4, UAS-GFP; Frt82B, atg13^Δ81^scrib^1^/Frt82B, tub-GAL80, atg13^Δ81^*

*y,w,ey-FLP/ey3.5-Hsp70-DmAtg13-GFP;UAS-Ras^V12^/act>y^+^>GAL4, UAS-GFP; Frt82B, atg13^Δ81^scrib^1^/Frt82B, tub-GAL80, atg13^Δ81^*

**3i,j:** *y,w,ey-FLP/y,w;UAS-Ras^V12^/act>y^+^>GAL4, UAS-GFP; Frt82B, atg13^Δ81^scrib^1^/Frt82B, tub-GAL80, atg13^Δ81^*

**Figure 4.**

**4a:** *y,w,ey-FLP/y,w;UAS-Ras^V12^/act>y^+^>GAL4, UAS-GFP; Frt82B/Frt82B, tub-GAL80 y,w,ey-FLP/y,w;UAS-Ras^V12^/act>y^+^>GAL4, UAS-GFP; Frt82B, scrib^1^/Frt82B, tub-GAL80*

**4b:** *y,w,ey-FLP/y,w;UAS-Ras^V12^/act>y^+^>GAL4, UAS-GFP; Frt82B, scrib^1^/Frt82B, tub-GAL80 y,w,ey-FLP/y,w;UAS-Ras^V12^/act>y^+^>GAL4, UAS-GFP; Frt82B, atg13^Δ81^scrib^1^/Frt82B, tub-GAL80, atg13^Δ81^*

**4c:** *y,w,ey-FLP/y,w;UAS-Ras^V12^/act>y^+^>GAL4, UAS-GFP; Frt82B/Frt82B, tub-GAL80 y,w,ey-FLP/y,w;UAS-Ras^V12^/act>y^+^>GAL4, UAS-GFP; Frt82B, scrib^1^/Frt82B, tub-GAL80 y,w,ey-FLP/y,w;UAS-Ras^V12^/act>y^+^>GAL4, UAS-GFP; Frt82B, atg13^Δ81^scnb^1^/Frt82B, tub-GAL80, atg13^Δ81^*

*hsflp;;Frt82B, atg13^Δ81^/ Frt82B, atg13^Δ81^*

*hsflp;;Frt82B / Frt82B*

**Figure 5.**

**5a:** *y,w,ey-FLP/y,w;UAS-Ras^V12^/act>y^+^>GAL4, UAS-GFP; Frt82B/Frt82B, tub-GAL80 y,w,ey-FLP/y,w;UAS-Ras^V12^/act>y^+^>GAL4, UAS-GFP; Frt82B, scrib^1^/Frt82B, tub-GAL80 y,w,ey-FLP/y,w;UAS-Ras^V12^/act>y^+^>GAL4, UAS-GFP; Frt82B, atg13^Δ81^scrib^1^/Frt82B, tub-GAL80, atg13^Δ81^*

**5b:** *y,w,ey-FLP/y,w;UAS-Ras^V12^/act>y^+^>GAL4, UAS-GFP; Frt82B/Frt82B, tub-GAL80*

**5c,d:** *y,w,ey-FLP/y,w;UAS-Ras^V12^/act>y^+^>GAL4, UAS-GFP; Frt82B, scrib^1^/Frt82B, tub-GAL80*

**5e:** *y,w,ey-FLP/y,w;UAS-Ras^V12^/act>y^+^>GAL4, UAS-GFP; Frt82B, atg13^Δ81^scrib^1^/Frt82B, tub-GAL80, atg13^Δ81^*

**5g:** *y,w,ey-FLP/y,w;UAS-Ras^V12^/act>y^+^>GAL4, UAS-GFP; Frt82B, scrib^1^/Frt82B, tub-GAL80*

**Expanded View Figure 1.**

**EVF 1a:** *y,w,ey-FLP/y,w;UAS-Ras^V12^/act>y^+^ >GAL4, UAS-GFP; Frt82B/Frt82B, tub-GAL80*

**EVF 1b,c:** *y,w,ey-FLP/y,w;UAS-Ras^V12^/act>y^+^ >GAL4, UAS-GFP; Frt82B, scrib^1^/Frt82B, tub-GAL80*

**EVF 1d:** *w^1118^*

**EVF 1e:** *w^1118^/w^*^, spok-Gal4, UAS-Dcr-2.D; UAS-rpr*

**EVF 1f:** *;;ecd^1^, st^1^, ca^1^/ecd^1^, st^1^, ca^1^* (Bloomington 218)

**Expanded View Figure 2.**

**EVF 2a:** *y,w,ey-FLP/y,w;UAS-Ras^V12^, pmCh-atg8a/act>y^+^>GAL4, UAS-GFP; Frt82B/Frt82B, tub-GAL80*

*y,w,ey-FLP/y,w;UAS-Ras^V12^, pmCh-atg8a/act>y^+^>GAL4, UAS-GFP; Frt82B, scrib^1^/Frt82B, tub-GAL80*

*y,w,ey-FLP/y,w;UAS-Ras^V12^, pmCh-atg8a/act>y^+^>GAL4, UAS-GFP; Frt82B, atg13^Δ81^ scrib^1^/Frt82B, tub-GAL80, atg13^Δ81^*

**EVF 2b:** *y,w,ey-FLP/y,w;UAS-Ras^V12^/act>y^+^ >GAL4, UAS-GFP; Frt82B, atg13^Δ81^ scrib^1^/Frt82B, tub-GAL80, atg13^Δ81^*

**EVF 2c:** *y,w,ey-FLP/y,w;UAS-Ras^V12^/act>y^+^>GAL4, UAS-GFP; Frt82B/Frt82B, tub-GAL80 y,w,ey-FLP/y,w;UAS-Ras^V12^/act>y^+^>GAL4, UAS-GFP; Frt82B, scrib^1^/Frt82B, tub-GAL80 y,w,ey-FLP/y,w;UAS-Ras^V12^/act>y^+^>GAL4, UAS-GFP; Frt82B, atg13^Δ81^ scrib^1^/Frt82B, tub-GAL80, atg13^Δ81^*

**EVF 2d,e:** *y,w,ey-FLP/y,w; g-atg13, UAS-Ras^V12^/act>y^+^>GAL4, UAS-GFP; Frt82B, atg13^Δ81^ scrib^1^/Frt82B, tub-GAL80, atg13^Δ81^*

**EVF 2f:** *y,w,ey-FLP/y,w;UAS-Ras^V12^/act>y^+^>GAL4, UAS-GFP; Frt82B, atg14^Δ5.2^, scrib^1^/Frt82B, tub-GAL80, atg14^Δ5.2^*

**EVF 2g:** *y,w;;Frt82B, atg13^Δ81^/Frt82B, atg13^Δ81^ y,w,ey-FLP/y,w;UAS-Ras^V12^/act>y^+^>GAL4, UAS-GFP; Frt82B, scrib^1^/Frt82B, tub-GAL80 y,w,ey-FLP/y,w;UAS-Ras^V12^/act>y^+^>GAL4, UAS-GFP; Frt82B, atg14^Δ5.2^, scrib^1^/Frt82B, tub-GAL80, atg14^Δ5.2^*

**Expanded View Figure 3.**

**EVF 3a:** *y,w,ey-FLP/y,w;UAS-Ras^V12^/act>y^+^>GAL4, UAS-GFP; Frt82B, atg13^Δ1^ scrib^1^/Frt82B, tub-GAL80, atg13^Δ81^*

*y,w,ey-FLP/y,w;UAS-Ras^V12^/act>y^+^>GAL4, UAS-GFP; Frt82B, scrib^1^/Frt82B, tub-GAL80*

**EVF 3b:** *y,w,ey-FLP/y,w;UAS-Ras^V12^/act>y^+^>GAL4, UAS-GFP; Frt82B, atg13^Δ1^ scrib^1^/Frt82B, tub-GAL80, atg13^Δ81^*

*y,w,ey-FLP/y,w;UAS-Ras^V12^/act>y^+^>GAL4, UAS-GFP; Frt82B/Frt82B, tub-GAL80*

**EVF 3c:** *y,w,ey-FLP/y,w;UAS-Ras^V12^/act>y^+^>GAL4, UAS-GFP; Frt82B/Frt82B, tub-GAL80 y,w,ey-FLP/y,w;UAS-Ras^V12^/act>y^+^>GAL4, UAS-GFP; Frt82B, scrib^1^/Frt82B, tub-GAL80 y,w,ey-FLP/y,w;UAS-Ras^V12^/act>y^+^>GAL4, UAS-GFP; Frt82B, atg13^Δ81^ scrib^1^/Frt82B, tub-GAL80, atg13^Δ81^*

**EVF 3d:** *y,w,ey-FLP/y,w;UAS-Ras^V12^/act>y^+^>GAL4, UAS-GFP; Frt82B/Frt82B, tub-GAL80 y,w,ey-FLP/y,w;UAS-Ras^V12^/act>y^+^>GAL4, UAS-GFP; Frt82B, scrib^1^/Frt82B, tub-GAL80 y,w,ey-FLP/y,w;UAS-Ras^V12^/act>y^+^>GAL4, UAS-GFP; Frt82B, atg13^Δ81^ scrib^1^/Frt82B, tub-GAL80, atg13^Δ81^*

**Expanded View Figure 4.**

**EVD 4a:** *y,w,ey-FLP/y,w;UAS-Ras^V12^/act>y^+^>GAL4, UAS-GFP; Frt82B, scrib^1^/Frt82B, tub-GAL80*

